# Acute supplementation of beta-hydroxybutyrate increases visual cortical excitability in humans: a combined EEG and MRS study

**DOI:** 10.1101/2025.03.11.642553

**Authors:** Cecilia Steinwurzel, Maria Concetta Morrone, Ele Ferrannini, Francesca Frijia, Domenico Montanaro, Giuseppe Daniele, Paola Binda

## Abstract

Increasing plasma levels of ketone bodies via supplementation or dieting has been repeatedly used to ameliorate neurological symptoms and enhance cognitive performance. Here we aim to gain insight into the underlying mechanisms by characterizing the acute effects on visual cortical function of a single dose β-hydroxybutyrate (βHB) supplementation. Young human volunteers were orally administered a βHB ester; we used EEG to assess cortical excitability and responsivity to visual stimulation, and Magnetic Resonance Spectroscopy to quantify glutamate and GABA+ concentrations in the occipital cortex. βHB supplementation increased the amplitude of Visual Evoked Potentials and enhanced the resting-state EEG alpha rhythm. These electrophysiological changes were paralleled by a neurometabolic change in the occipital cortex, where glutamate (but not GABA+) concentration increased. The glutamate increase was correlated with the increased Visual Evoked Potential amplitude. This suggests that acute βHB supplementation increases the excitability of the brain cortex, as assessed neurometabolically and electrophysiologically.

## Introduction

Brain metabolism primarily relies on glucose (Owen *et al*., 1967). However, during fasting and strenuous physical exertion, ketone bodies serve as the primary alternative source of energy for brain cells (Jensen *et al*., 2020). Ketogenesis predominantly occurs in hepatocytes, where free fatty acids undergo a process of β-oxidation regulated by insulin and glucagon (Puchalska & Crawford, 2017). The three final products, acetone, acetoacetate and β-hydroxybutyrate (βHB) (Newman & Verdin, 2017), are released into the blood stream and they cross the blood-brain barrier via monocarboxylic acid transporters (Jensen *et al*., 2020).

The level of ketone bodies in the bloodstream can also be elevated through exogenous supplementation of ketone esters or salts and through ketogenic diets (Clarke *et al*., 2012; Stubbs *et al*., 2017; Soto-Mota *et al*., 2020). Even a single-dose of acute βHB supplementation produces a substantial raise of βHB levels in the blood stream and increased βHB concentrations in the brain, as estimated with Magnetic Resonance Spectroscopy (^1^H-MRS) (Mujica-Parodi *et al*., 2020; Hone-Blanchet *et al*., 2023). Using MRS with ^13^C labelling also provided evidence that peripherally infused βHB is metabolized in brain cells, particularly neurons, as the ^13^C label was detectable in the amino acid pools of glutamate, glutamine, and aspartate (Pan *et al*., 2002).

The exploration of how the levels of ketone bodies could impact brain function dates back to the 1920s, when a low-carbohydrate/high-fat diet demonstrated efficacy in the treatment of pediatric epilepsy (Helmholz & Keith, 1933). Although the use of ketogenic diets declined with the advent of anticonvulsant drugs, interest resurfaced in the past two-three decades, due to the ability of these diets to reduce seizures in drug-resistant epilepsy (Thiele, 2003; D’Andrea Meira *et al*., 2019; Poff *et al*., 2019) and their association with cognitive enhancement, observed in diverse human populations (from young healthy participants to elderly patients, with or without cognitive impairment) and animal models (Chinna-Meyyappan *et al*., 2023). Many studies aimed to unravel the complex effects of ketone bodies on neuronal function (Achanta & Rae, 2017). Invitro and ex-vivo studies outlined a number of pathways through which βHB could modulate neuronal function, including: 1) by modulating the energy balance and expenditure in the cell, 2) by acting as extracellular receptor ligand, 3) by influencing intracellular signaling post-translationally (for review, Garcia-Rodriguez & Gimenez-Cassina, 2021), 4) by acting on the synthesis and release of glutamate (Juge *et al*., 2010) and GABA (Erecinska *et al*., 1996; Yudkoff *et al*., 2007; Yudkoff *et al*., 2008).

A recent human study with 7T Magnetic Resonance Spectroscopy (^1^H-MRS) (Hone-Blanchet *et al*., 2023) provided partial support for the latter pathway, showing that acute βHB supplementation reduces GABA levels by about 50% (Maddock, 2023) within the cingulate cortices, paralleling the increase of βHB concentration in the same voxel. Intriguingly, the GABA decrease was accompanied by a glutamate decrease and the decrease of both GABA and glutamate was mainly observed in older participants, while no change or even a slight increase was detectable for the youngest participants in the sample (around 25 years old). These features make it hard to predict whether and how these large but coordinated GABA and glutamate concentration changes could influence neuronal activity and function.

One established methodology to non-invasively assess the activity and function of the brain cortex is through electro-encephalography (EEG), either performed in combination with visual stimulation or in resting-state conditions. The resting-state EEG power spectrum indexes endogenous rhythms and thereby informs about the dynamics of the recurrent connectivity of the brain. One main index of interest is the power in the alpha band within the occipital cortex, which is thought to be related to cortical excitability (Lozano-Soldevilla, 2018). A visual stimulus that oscillates in time entrains a cortical rhythm readily measured with EEG and termed steady-state Visual Evoked Potential (VEP). VEP amplitude provides a sensitive index of visual cortex excitability (Hudnell & Boyes, 1991), indicative of the glutamate to GABA balance (Boker & Heinze, 1984; Bartel *et al*., 1988; Bale *et al*., 2005; Geller *et al*., 2005). In addition, by overlaying visual stimuli of different orientations (cross-orientation stimuli), steady-state Visual Evoked Potentials reveal a strong reciprocal inhibition, previously shown to depend on GABAergic signaling (Morrone *et al*., 1982; Morrone & Burr, 1986; Morrone *et al*., 1987; Smith *et al*., 2006).

Here we used resting-state EEG and steady-state VEPs to evaluate the acute effects of a single dose supplementation of a βHB ester. In the same session, we also used MEGA-PRESS MRS to measure GABA+ and glutamate levels in the occipital (visual) cortex. To preview our results, we find increased cortical excitability with EEG, correlated with increased occipital glutamate levels, and no associated modulation of GABA+.

## Materials and methods

### Experimental model and subject details

Experimental procedures were approved by the regional ethics committee [*Comitato Etico Regionale per la sperimentazione clinica della Regione Toscana — Sezione Area Vasta Nord Ovest*; protocol “Studio BetaidrossiBrain: effetto del betaidrossibutirrato sulla attivita’ della corteccia cerebrale”] and are in line with the declaration of Helsinki. Participants gave written informed consent prior to inclusion and completed a confidential medical screening questionnaire to determine eligibility.

We recruited 12 healthy volunteers (2 females, 10 males). The selection criteria included normal or corrected-to-normal vision, no known history of amblyopia, eye surgery, or other active eye disease. One female participant was excluded due to an anatomical abnormality encountered in the MR anatomical acquisition. One male participant was discarded due to an abnormal fasting glycemia (>140 mg/dl). Participants came to the CNR-Fondazione Regione Toscana G. Monasterio (Pisa, Italy) at 8.00 a.m, after an overnight fast of 12 hours. Blood samples were collected upon arrival, followed by the MRI (Magnetic Resonance Imaging) acquisitions and the EEG recordings. These measurements were all repeated after the βHB supplementation. This was achieved by orally administering a single dose of a βHB ester (HVMN™), containing 25 grams of D-β-hydroxybutyrate. One hour later, EEG recordings were repeated and, at about 2 hours after βHB supplementation, the second MRI acquisition was performed. The whole experimental procedure could last up to 8 hours per participant, which limited the sample size we could achieve. However, a power analysis based on a recent study measuring the impact of βHB supplementation upon neurometabolites concentrations through MRS (Hone-Blanchet *et al*., 2023), indicates that a sample size of 9 participants would be sufficient to detect the large observed modulation of Glutamate concentrations (Cohen’s d = 1.13, computed from their Figure 3B) given an a priori chose alpha level of 0.05 (two-tailed) and a power of 0.80.

## Methods details

### Collection and analysis of blood samples

Venous blood samples for pharmacokinetic analysis were collected before starting the experimental procedure and at 60, 120, and 180 min after βHB supplementation (Figure 1A). Blood samples were stored on ice, centrifuged (3000 rpm for15 min at 4 °C) and duplicate plasma aliquots were stored at −80 °C. Plasma concentrations of βHB, glucose, insulin, C-peptide, glucagon, GLP-1 total and active were assessed for each time-point, using the following procedures. βHB was stored in a K2E (EDTA) 7.2 mg BD Vacutainer test tube and assayed using the commercially available ß-hydroxybutyrate LiquiColor® (EKF Diagnostics). Plasma glucose was stored in an FX 10mg 8mg, BD Vacutainer and was assessed using a commercial Glucometer (YSI 2300 STAT Plus). Insulin and C-peptide were stored in Litio/Heparina 102 IU BD Vacutainer and measured using a commercially available ELISA assay (Mercodia, DBA Italia S.r.l.). Glucagon, GLP-1 total and active were stored using a BD™ P800 Blood Collection System test tube. Glucagon was assayed using a commercially available ELISA assay (Mercodia, DBA Italia S.r.l.), whilst GLP-1 levels were assessed with dedicated commercially available ELISA kit (Merck Life Science S.r.l.).

**Figure 1.**
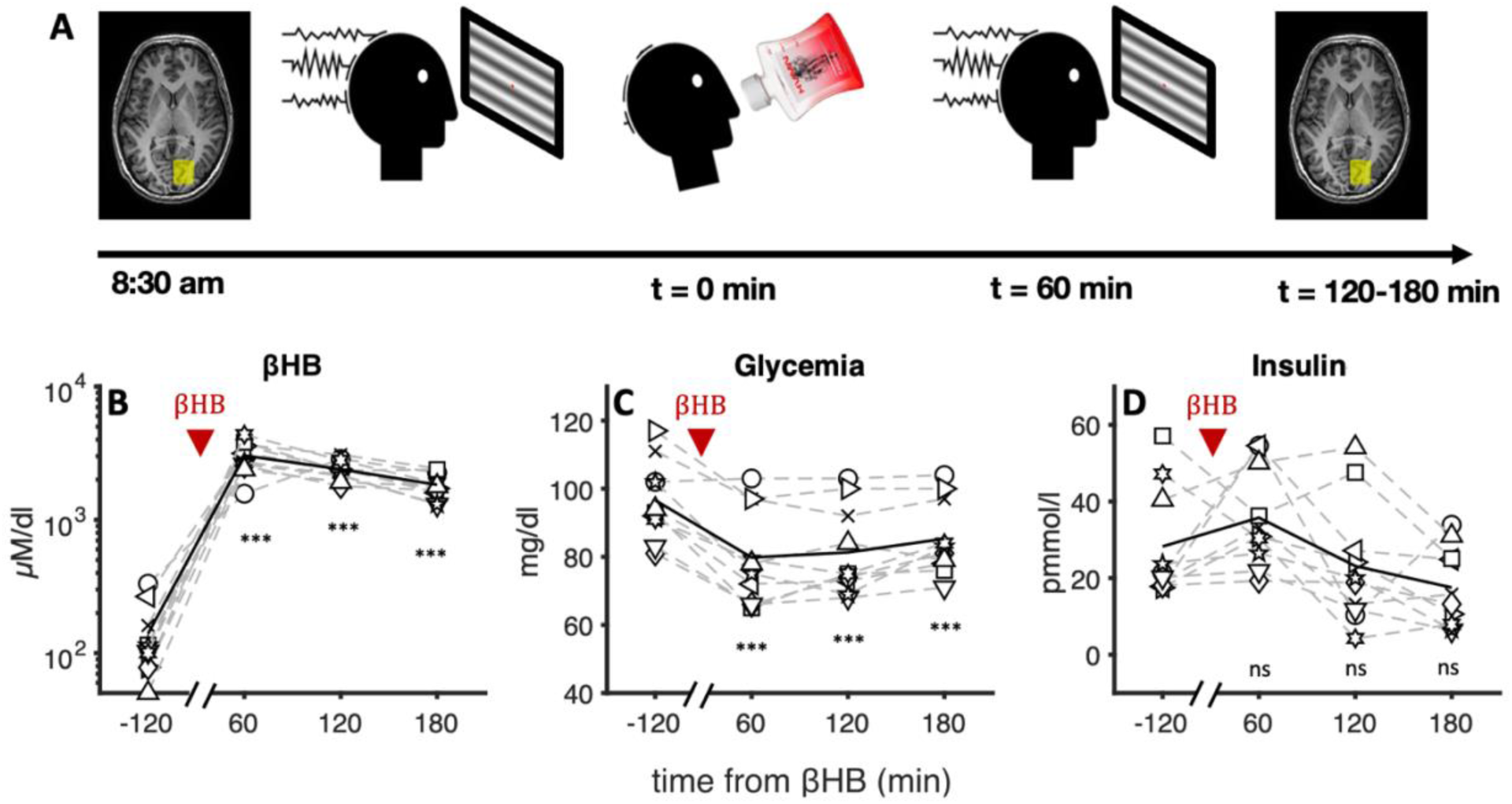
Experimental design and time-courses of metabolic parameters. A) Experimental procedure. The experiment lasted about 6 hours per participant, with metabolic, EEG and MRS measurements taken at baseline and after βHB supplementation. B) Plasma concentrations of βHB C) Glucose D) Insulin, at baseline and 60, 120 and 180 minutes after βHB supplementation (marked by red triangle). In each panel, the solid black line shows the average time-course across participants, with individual participants shown as open symbols (symbol-participant pairing is consistent across all figures). Text insets give the significance of post-hoc tests comparing post-supplementation measures with baseline (ns, nonsignificant, *p < 0.05, **p < 0.01, ***p < 0.001, Bonferroni corrected).

### EEG recordings

EEG was recorded using a wireless g.Nautilus system, with a sampling rate of 250 Hz. The scalp electrodes were positioned according to the 10-20 international system and the reference electrode was positioned on the right earlobe. The impedance was checked before each recording and kept below 50 kΩ. For the present study, we recorded from 8 electrodes, including central electrodes *FZ*, *CZ* and the posterior and occipital electrodes *PZ*, *OZ*, *PO3*, *PO7*, *PO4*, *PO8*.

Visual stimuli were generated with Psychtoolbox for Matlab (Matlab r2017b, The Mathworks, Inc.) and displayed on a gamma-calibrated Barco monitor (Barco CDCT 6551, 800 × 600 pixels, 100 Hz) through a ViSaGe (CRS, Cambridge Research Systems, Rochester, UK). A custom-made trigger connected the ViSaGe with the g.Nautilus system. Participants were positioned at 114 cm from the screen. Two types of steady-state Visual Evoked Potentials (VEP) were tested; in addition, EEG recordings in resting state conditions were acquired while participants kept their eyes open and fixated on fixation spot on an otherwise empty monitor screen.

For steady-state VEP, the stimulus was a whole-screen horizontal sinusoidal grating (SF of 1 c/°, Michaelson contrast of 35%, mean luminance of 51.8 cd/m^2^), modulated over time according to a sinusoidal function with a frequency of 8.33 Hz. In a separate recording, we also measured Cross-Orientation Inhibition (Morrone & Burr, 1986), by superimposing the same test stimulus with a vertical grating mask (SF of 0.8 c/°, Michaelson contrast of 45%) modulated sinusoidally at a frequency of 7.1 Hz. In this condition, the response to the test is strongly attenuated and the final response is proportional to the product of excitation by inhibition (Carandini & Heeger, 2011).

### MRI and MRS acquisitions

We acquired MRI data on a 3-T scanner (GE HDx TWINSPEE, GE Medical Systems, Wisconsin, Milwaukee, USA) at CNR-Regione Toscana G. Monasterio (Pisa, Italy) using an 8-channel head coil equipped with TwinSpeed gradients.

We collected a three-dimensional (3D) fast spoiled gradient recall T1-weighted images (*3D SPGR*, TR/TE = 10.7/4.9 ms, FOV = 25.6 cm, acquisition matrix = 256×256, voxel size = 1mm isotropic, slice gap = 0mm, BW = 15.6kHz, NEX = 1) covering the entire brain. We collected MRS data with a MEGA-PRESS (Mescher *et al*., 1998) sequence (TE = 68 ms; TR = 1500 ms; 320 transients of 4,096 data points acquired in 8-min experiment time; 16-ms Gaussian editing pulse applied at 1.9 (ON) and 7.46 (OFF) ppm); for further information, please see Supplemental Table S1, which was set-up following consensus guidelines (Lin *et al*., 2021).

We acquired spectra from two MRS voxels (25 × 25 × 25 mm^3^): before and after βHB supplementation. We manually positioned the voxel based on each participant’s anatomical T1 scan, ensuring to maintain the same voxel placement before and after βHB supplementation. To cover visual areas, we centered the MRS voxel on right the calcarine sulcus (Figure 3A) in all but two participants, where the voxel was centered on the left calcarine sulcus to minimize inclusion of the ventricular space. During the MRS acquisitions participants kept their eyes closed.

### Quantification and statistical analysis

#### EEG analysis

Offline analyses were performed using EEGlab (14.1.2b) and custom-made MATLAB scripts (MATLAB 2019a; MathWorks). For Visual Evoked Potentials, the EEG signal was divided into epochs of 240 ms (2 cycles of stimulation), synchronized with the alternation of the grating stimulus. The raw EEG signal was visually inspected and epochs with obvious artifacts were discarded from further analyses. Valid epochs were averaged (separately for each participant and condition, pre vs. post βHB supplementation) and a Fast Fourier Transform (FFT) was applied to the average trace. The amplitude of the second harmonic (twice the stimulus frequency, i.e. 16.6 Hz) was used to index the visually evoked response.

The resting state EEG signal was divided into 10-second-long epochs, which were visually inspected and to reject epochs with obvious artifacts. A FFT was then applied to each epoch, and the amplitude spectrum was averaged across epochs (separately for each participant and condition). The power in the alpha frequency range (8-13 Hz) was integrated and then square-rooted to obtain alpha-amplitude.

#### MRS analysis

MRS data pre-processing was performed using MRSpa v1.5 (https://www.cmrr.umn.edu/downloads/mrspa/). Eddy current, frequency, and phase correction were applied before subtracting the average ON and OFF spectra, resulting in one edited spectrum per participant and condition. We used LCModel (Provencher, 2001), with dedicated MEGA-PRESS basis set (‘mega-press-3’) to fit the spectra within the chemical shift range 1.9 to 4 ppm (Figure 3B) to avoid artifacts contamination, as suggested by the LCModel Manual (Provencher, 2001). We estimated the concentration of the following neurometabolites: glutamate (Glu), glutamine (Gln), γ-aminobutyric acid (GABA+), glutathione (GSH), and *N*-acetylaspartate (NAA). We refer to GABA concentration as ‘GABA+’, as MRS measurements of GABA with MEGA-PRESS are known to include coedited macromolecules (Mullins *et al*., 2014).

We focused on glutamate rather than GLX as βHB has been shown to directly affect glutamate’s levels (Juge *et al*., 2010; Hone-Blanchet *et al*., 2023). Previous work using MEGA-PRESS at 3T has shown that glutamate and glutamine signal can be separable and reliably quantified using LCModel (Sanaei Nezhad *et al*., 2018). Our glutamate measurements are in line with the spectral quality criteria outlined in previous work (Sanaei Nezhad *et al*., 2018)(Jia *et al*., 2022), and both glutamate and GABA + had Cramér–Rao lower bound (CRLB) values smaller than 10%. Given that it cannot be taken for granted that βHB supplementation leaves other metabolites like NAA unaffected, we elected to reference our GABA+ and glutamate concentrations to water. To ensure that our results were not driven by variability in voxel tissue composition, metabolite concentration values were further corrected using the alpha-method described by Harris et al (Harris *et al*., 2015):

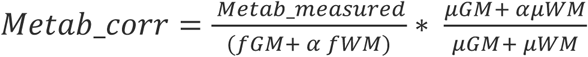

Where fGM and fWM are the fractions of individual gray and white matter content in each participants voxel, μ_GM_ and μ_WM_ are the GM and WM fractions of the group average voxel fractions, and alpha value is set to 0.5 (Harris *et al*., 2015).

CSF, gray (fGM) and white matter (fWM) fractions were extracted through co-registering high resolution three-dimensional T1 images (segmented with FreeSurfer (Fischl *et al*., 2002)) with a binary MRS voxel mask. There was no significant difference between CSF (fCSF_pre_ = 5.2%, fCSF_post_= 5.5%, t(9) = 0.4), gray (fGM_pre_ = 47.9 %, fGM_post_= 48.4 %, t(9) = 0.8), and white (fWM_pre_ = 47.2 %, fWM_post_= 45.9 %, t(9) = 0.6), matters segmentation values between baseline and post βHB supplementation.

## Results

### Acute βHB supplementation increases βHB levels in the bloodstream

Before and after supplementation of a βHB ester, we monitored the main metabolic parameters through hourly blood sampling for 3 hours (Table 1).

**Table 1.** Anthropometrics and metabolic variables, reported as mean (SD)

| Variables | Baseline | 60 min | 120 min | 180 min |
| --- | --- | --- | --- | --- |
| Age (years) | 27 (4) | - | - | - |
| BMI (kg/m<sup>2</sup>) | 22.4 (2.2) | - | - | - |
| βHB (μmol/L) | 140 (90) | 3036 (826) | 2393 (415) | 1834 (408) |
| Glucose (mg/dL) | 97 (12) | 80 (14) | 81 (13) | 85 (11) |
| Insulin (μU/mL) | 24 (14) | 36 (13) | 23 (16) | 18 (10) |
| C-peptide (ng/mL) | 1.2 (0.5) | 1.3 (0.5) | 1.1 (0.4) | 0.8 (0.4) |
| Glucagon (ng/mL) | 7.0 (4.0) | 9.7 (10.1) | 10.8 (9.8) | - |
| GLP1-total (ng/mL) | 37.9 (18.9) | 38.6 (24.4) | 40.8 (23.4) | - |
| GLP1-active (ng/mL) | 4.8 (1.7) | 2.5 (2.4) | 3.7 (2.5) | - |

Plasma βHB concentrations (Figure 1B) rose from a mean baseline level of 140 µmol/L to a maximum of 3036 µmol/L one hour after βHB supplementation, in line with previous reports (Clarke *et al*., 2012; Mikkelsen *et al*., 2015; Stubbs *et al*., 2017). The variation was significant (one-way ANOVA for repeated measures, F(3,27) = 74.3, p < 0.001). After the peak response, plasma βHB decreased non-linearly(Stubbs *et al*., 2017), and was still significantly elevated 180 min following ingestion (post-hoc t = 8.31, Bonferroni corrected p < 0.001).

Plasma glucose levels (Figure 1C) decreased with time (F(3,27) = 24.8, p < 0.001) and remained below baseline throughout the experiment (all post-hoc t > 5.2, all Bonferroni corrected p < 0.001).

Plasma insulin concentrations (Figure 1D) showed a subtler and more variable change (F(3,27) = 5.3, p = 0.005), with a non-significant rise at 60 min after βHB supplementation (post-hoc t = 1.59, Bonferroni corrected p = 0.755), followed by a progressive non-significant decreasing trend (all post-hoc t < 2.2, all Bonferroni corrected p > 0.197). The plasma C-peptide levels showed a similar trend as insulin levels.

The active fraction of GLP1 was significantly modulated (F(2,18) = 4.1, p = 0.034), with a decrease at 60 min (post-hoc t = 2.87, Bonferroni corrected p = 0.030). The other parameters (GLP1-total and glucagon) did not show reliable changes (both F(2,18) < 1.45, both p > 0.261 after correction with Greenhouse-Geisser).

### Acute βHB supplementation increases alpha and VEP amplitude

To investigate whether βHB modulates brain activity, we used EEG to measure endogenous rhythms in resting state (with eyes open) and Visual Evoked Potentials. We acquired these measures immediately before βHB supplementation (blue lines in Figure 2) and 60 minutes afterwards (red lines), corresponding to the peak βHB plasma level.

**Figure 2.**
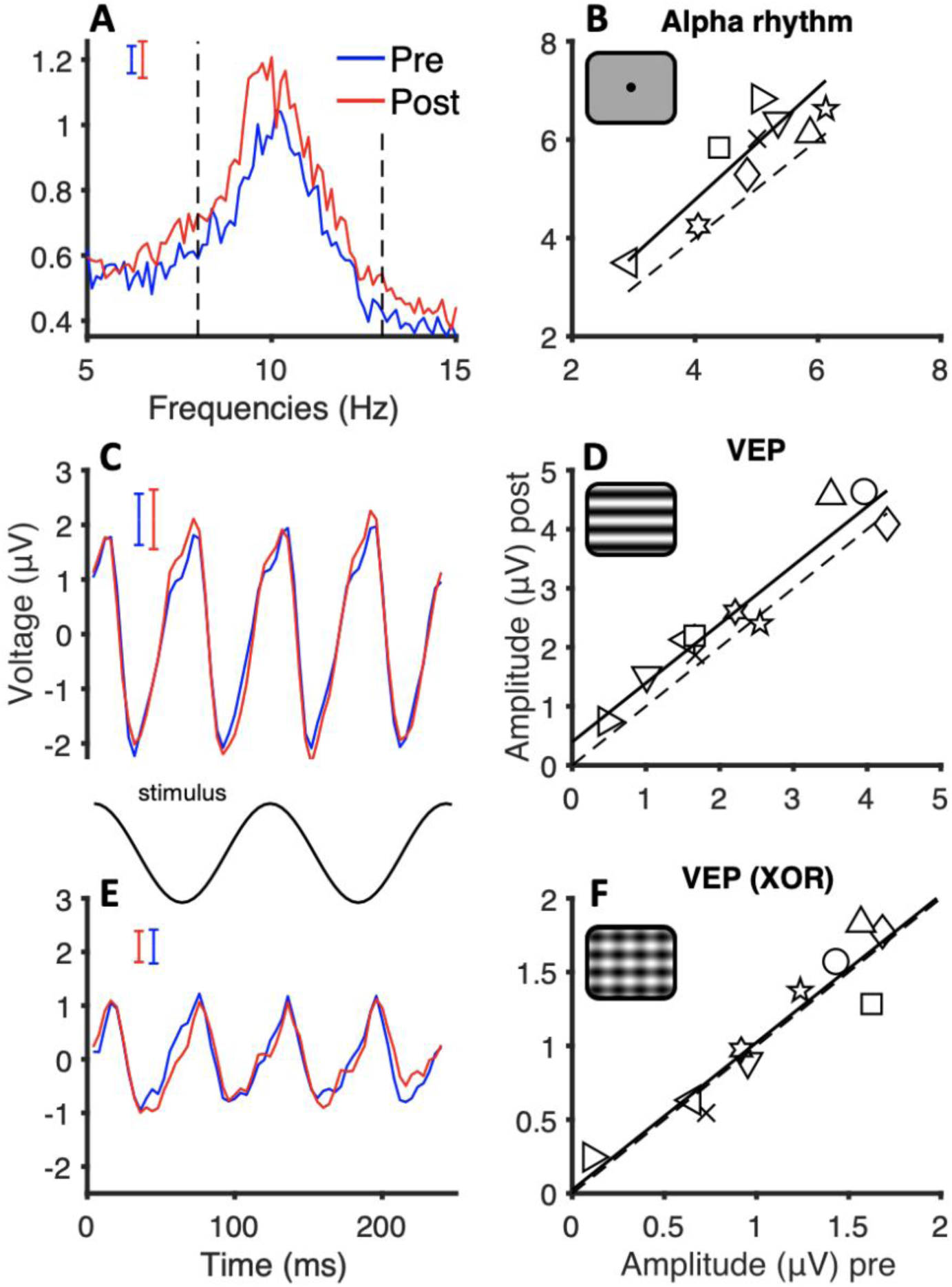
Effects of βHB supplementation on EEG recordings. A) Amplitude spectra of EEG recordings measured in resting-state with eyes open, immediately before (blue) and 60 minutes after (red) βHB supplementation; the vertical dashed lines mark the frequency range used to extract the amplitude of the alpha rhythm (8-13Hz). B) Alpha amplitude in each participant, values acquired after βHB supplementation plotted against the corresponding values acquired before it. All data points are above the bisection of the axes (dashed line), implying that alpha amplitude was consistently increased. C) Waveforms of steady-state Visual Evoked Potentials, elicited by horizontal grating oscillating in time at 8.33 Hz. D) VEP amplitude (second harmonic of the stimulus frequency), post-vs. pre-supplementation, again showing a consistent enhancement in most participants. E) Waveforms of steady-state Visual Evoked Potentials, elicited by combining the same horizontal grating oscillating in time at 8.33 Hz with an orthogonal grating oscillating at 7.1 Hz, yielding markedly smaller responses (compare with panel C), as expected from cross-orientation inhibition (XOR). F) VEP-XOR amplitude, showing no change after βHB supplementation. In panels A, C and E, curves are averages across participants and the floating error bars show the s.e.m.. In panels B, D and F, the dashed line marks the y=x function and the continuous black line gives the best fitting linear function across the data points; icons in the top-left corners represent the visual stimulation conditions: no stimulus in B, a single horizontal grating in D and the same horizontal grating overlaid with a vertical grating in F.

Figure 2A shows the amplitude spectrum of the resting state EEG for the occipital electrode *OZ*. There is a marked amplitude increase within the alpha band (8-13 Hz, indicated by vertical dashed lines) after βHB supplementation, suggesting an overall increase of cortical excitability. Figure 2B shows that the alpha-amplitude (quantified as integral of amplitude values over the alpha band) was tightly correlated pre- and post-supplementation (Pearson’s r = 0.88 and p < 0.001), supporting the reliability of the measurements. The post-supplementation amplitudes were reliably higher than baseline (t(9) = 4.62, p < 0.001). A similar effect was seen across several other electrodes (*PO3*: t(9) = 4.01, p = 0.003; *PO4*: t(9) = 3.20, p = 0.011; *PO8*: t(9) = 3.51, p = 0.007; *PZ*: t(8) = 2.76, p = 0.025; *CZ*: t(9) = 3.58, p = 0.006; *FZ*: t(9) = 3.31, p = 0.011; across all these electrodes, pre- and post-supplementation alpha-amplitudes were correlated: all Pearson’s r > 0.93 and p < 0.001).

Figure 2C-F show the Visual Evoked Potentials for the occipital electrode *OZ*. Figure 2C displays the voltage modulation elicited by an alternating sinusoidal grating; as expected, this is mainly at the second harmonic (16.6 Hz) of the stimulus temporal frequency (8.3 Hz). Figure 2E shows the voltage modulation elicited by the same grating, coupled with an orthogonal mask alternating at a different frequency (7.1 Hz) that strongly attenuates the response – in line with previous cross-orientation inhibition studies (Morrone *et al*., 1982; Morrone & Burr, 1986; Morrone *et al*., 1987). Figures 2D and F report the amplitude of the second harmonic for both stimulus types, pre- and post-supplementation, again tightly correlated and supporting the reliability of the measurements (grating alone: Pearson’s r = 0.96, p < 0.001; grating plus mask: Pearson’s r = 0.94, p < 0.001). We evaluated the impact of βHB supplementation on the amplitude of responses to both stimulus types, with a two-way ANOVA for repeated measures with factors: time (pre- and post-supplementation) and stimulus (grating alone and grating plus orthogonal mask). This revealed a significant main effect of time (F(1,9) = 9.28, p = 0.014) and stimulus (F(1,9) = 24.38, p < 0.001), and a significant interaction between factors (F(1,9) = 8.36, p = 0.018). Post-hoc t-tests showed that βHB supplementation selectively affected Visual Evoked Potentials elicited by the individual grating (t = 4.2, Bonferroni corrected p = 0.003), while leaving responses to the grating plus orthogonal mask unaffected (t = 0.25, Bonferroni corrected p = 1). The observed effect of βHB supplementation on Visual Evoked Potentials elicited by the individual grating is large (Cohen’s d = 1.04), as the increment is an average of 17% of the amplitude before βHB supplementation.

### Acute βHB supplementation increases glutamate in occipital cortex

We used a dedicated MRS MEGA-PRESS sequence to evaluate GABA+ and glutamate concentrations within a 2.5 × 2.5 × 2.5 cm voxel placed in the occipital cortex (Figure 3A). Figure 3B shows the average MR Spectra (across participants) before βHB supplementation and an average of 120 minutes after it (blue and red lines respectively). We fit the spectra using LCModel (Provencher, 2001) to estimate the concentrations of the neurometabolites of interest (Table S1).

**Figure 3.**
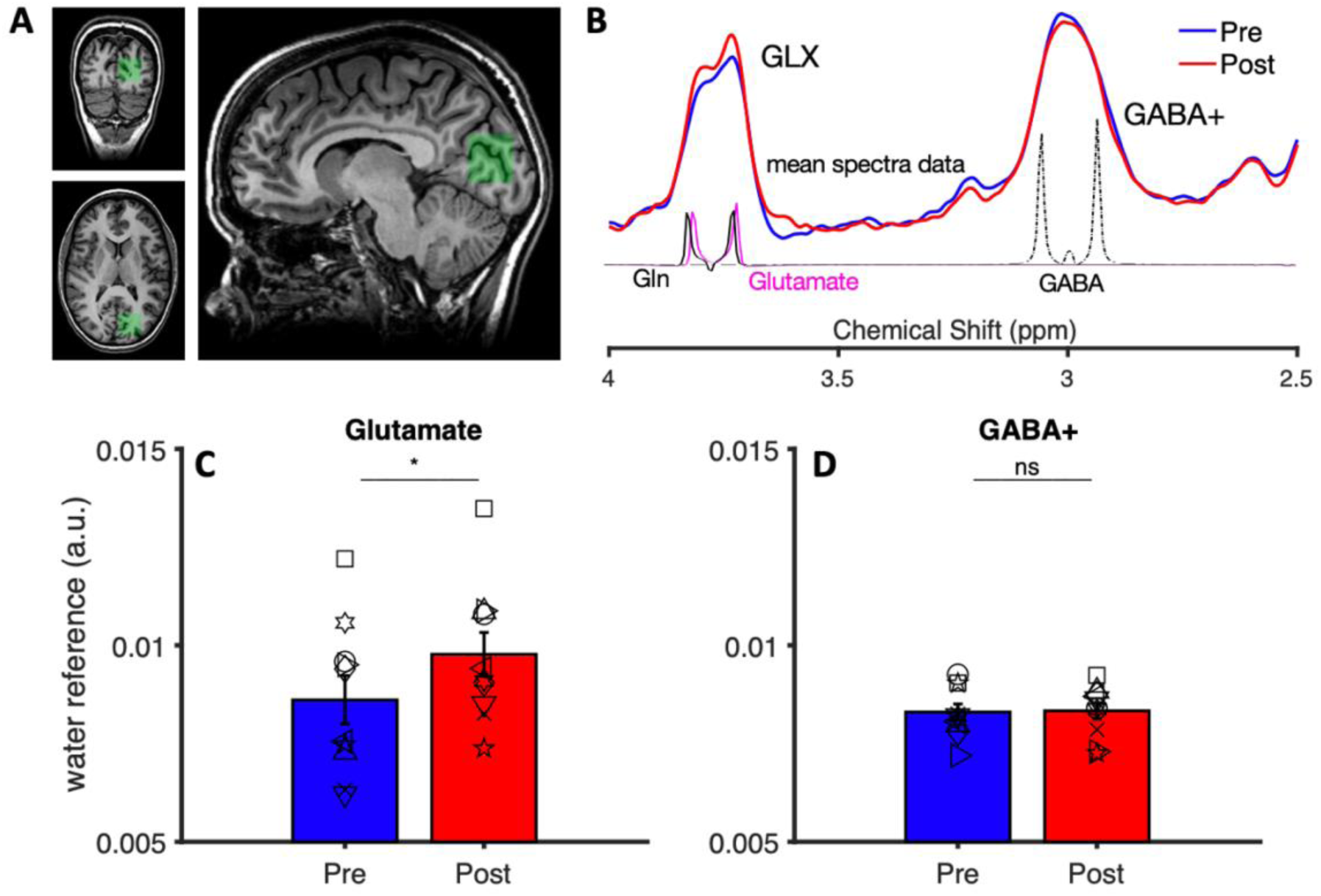
Effects of βHB supplementation on MEGA-PRESS MRS estimates of neurometabolite concentrations in the occipital cortex. A) Example of MRS voxel positioning for one example participant. B) MEGA-PRESS MRS spectra averaged across participants, acquired before and after βHB supplementation (blue and red, respectively), with the GLX and GABA+ peaks marked. The lower traces illustrate the basis functions used by LCModel to model the contribution of GABA+ (dotted black line), glutamate (Glu, pink solid line) and glutamine (Gln, solid black line) to the measured spectra. C-D) Estimated concentrations of glutamate and GABA+ (referenced to water). In all plots, open symbols show individual participants and error bars report s.e.m.. Text insets give the significance of post-hoc t-tests comparing post-supplementation measures with baseline (ns, nonsignificant, *p < 0.05).

After βHB supplementation, there was an increase of glutamate concentration (Figure 3C, Glu/water, t(9) = 2.52, p = 0.033) with no change in GABA+ (Figure 3D, GABA+/water, t(9) = 0.11 p = 0.91) or glutamine levels (Gln/water, t(9) = −1.16, p=0.274, *not shown*). A two-way ANOVA for repeated measures with factors time (pre and post-supplementation) and neurometabolite (glutamate and GABA+) revealed a significant time by neurometabolite interaction (F(1,9) = 5.88, p = 0.038), with no significant main effects (time: F(1,9) = 4.03, p = 0.075; neurometabolite: (1,9) = 3.04, p = 0.115). The increase of glutamate levels in the face of unaltered GABA+ levels is also seen as a significant increase of the glutamate to GABA+ ratio (t(9) = 2.32, p= 0.046, *not shown*); this ratio has been interpreted as an index of excitation/inhibition balance in the cortex (Steel *et al*., 2020; Rideaux *et al*., 2022), although this issue is controversial (Rideaux, 2021). Note that the glutamate concentration change is unlikely to result from changes in spectral fitting quality, as glutamate concentration levels estimated before and after βHB supplementation were correlated (r(10) = 0.70, p = 0.025, *not shown*). The glutamate increase is 13% of the glutamate levels before βHB supplementation (Cohen’s d = 0.8).

### The increases of glutamate concentration and VEP amplitude following acute βHB supplementation are correlated

We found a significant correlation between the increase of glutamate concentration measured with MRS (referenced on water and alpha corrected) and the increase of the visual evoked response measured with EEG (grating alone), as shown in Figure 4A (Pearson’s r = 0.67, p = 0.035). Interestingly, the levels of glutamate measured after βHB supplementation (Glu/water, alpha-corrected value) correlated with the rate of change in βHB plasma levels between 180 and 120 minutes (Figure 4B, Pearson’s r = 0.69, p = 0.026), which is the time window where spectra were acquired. We also explored correlations between MRS parameters, and the other significant change measured with EEG, the increased alpha-amplitude in resting state. However, no significant correlation emerged with either glutamate or GABA+ concentration changes (all |r| < 0.4, all p > 0.25).

**Figure 4.**
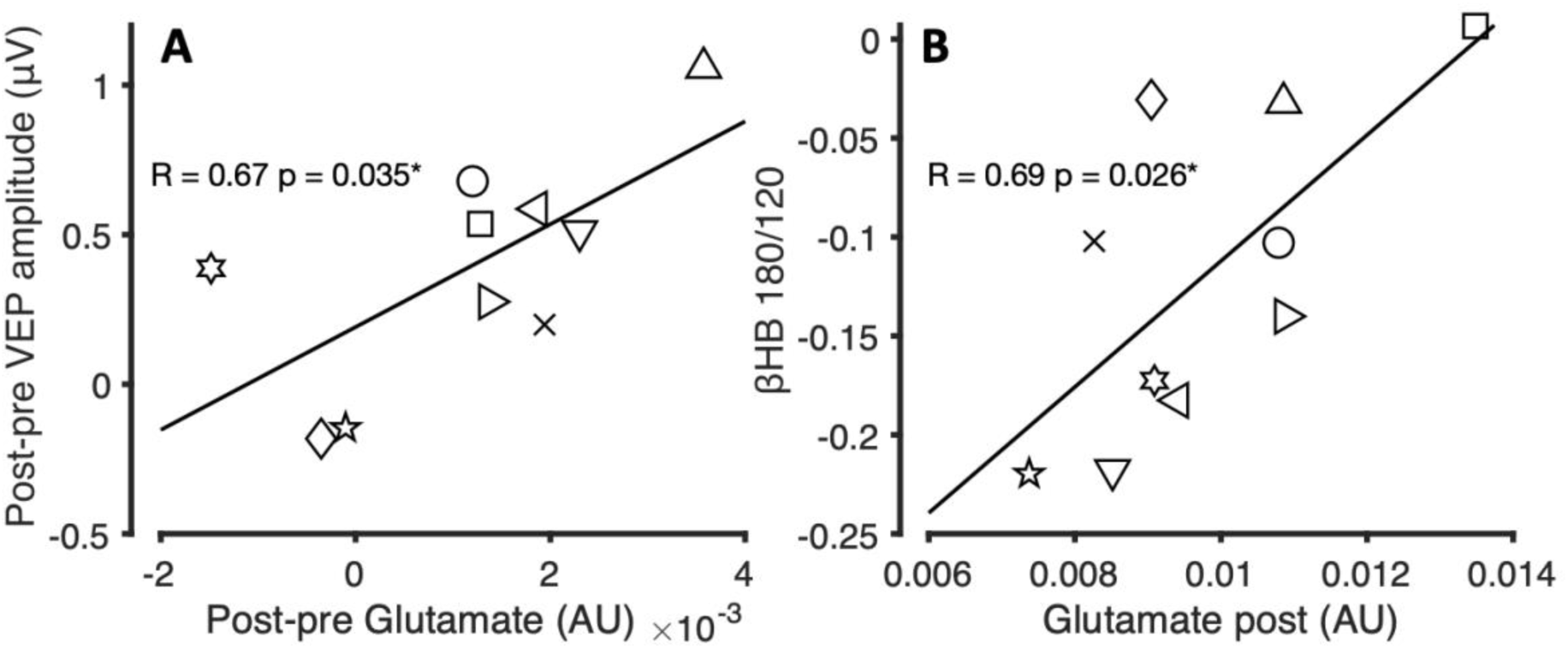
Correlations between MRS, EEG and βHB plasma levels. A) The amplitude change of Visual Evoked Potentials is directly correlated with the change of glutamate concentration (referenced to water) post-pre βHB supplementation. B) Glutamate concentrations after βHB supplementation (referenced to water) are directly with the rate of βHB clearance, measured as the ratio between βHB plasma levels at 2 and 3 hours after the supplementation. In both panels, the continuous black line gives the best fitting linear function across the individual participants data (open symbols); text insets give the Pearson’s correlation coefficients with associated p-value (*p < 0.05).

## Discussion

βHB supplementation in healthy young participants increased visual cortex excitability, as assessed by two independent EEG-physiological indices: steady-state Visual Evoked Potentials and resting-state alpha rhythm, suggestive of an increased ratio of excitation to inhibition in the visual cortex. This is supported by our MEGA-PRESS MRS results, showing increased glutamate concentrations and unaltered GABA+ concentrations in the occipital cortex (hence, increased glutamate to GABA+ ratio). Importantly, the increase of glutamate concentration was correlated with the enhanced Visual Evoked Potentials.

The resting-state alpha rhythm is the most prominent EEG pattern recorded from the occipital and parietal cortex; its amplitude is reliably modulated through pharmacological manipulation of the neural excitation/inhibition balance. Drugs known to potentiate GABAergic inhibition, like benzodiazepines, barbiturates, or anesthetics (Lozano-Soldevilla, 2018), lead to a consistent decrease of alpha power measured over parieto-occipital electrodes during resting-state. A similar alpha reduction can be obtained by suppressing glutamatergic transmission through blockade of glutamatergic receptors AMPA or NMDA (van Kerkoerle *et al*., 2014). In line with this pharmacological evidence showing that alpha amplitude can be modulated by the relative strength of glutamatergic versus GABAergic signals, we find that βHB supplementation produced a widespread increase of alpha amplitude, which accompanied an acute increase of glutamate (and unaltered GABA+) levels in the occipital cortex.

The amplitude of steady-state Visual Evoked Potentials has also been reliably linked with the balance between excitatory and inhibitory cortical signaling, as VEP amplitude is enhanced when the glutamate to GABA ratio is increased (Hudnell & Boyes, 1991; Bale *et al*., 2005; Geller *et al*., 2005). Coherently, we observed increased VEPs paralleling the increased concentration of occipital glutamate levels. In contrast, we found that the (attenuated) VEP responses observed with the superposition of an orthogonal mask (cross-orientation inhibition stimuli) were unaffected by βHB supplementation. This observation can be explained assuming that βHB supplementation enhanced both the grating and the mask response in a similar way (i.e. enhancing the excitatory drive to the cortex); given the divisive nature of the cross-orientation inhibition phenomenon (Morrone & Burr, 1986; Carandini & Heeger, 2011; Atallah *et al*., 2012), this is expected to result unaltered responses to the composite stimulus, as observed. Thus, the results from our cross-orientation inhibition stimuli may be interpreted as indirect evidence that βHB supplementation did not alter GABAergic signaling, in line with our MRS based GABA concentration estimates.

Although several previous studies investigated the effects of ketogenic diets or ketone supplementation on brain function, none simultaneously monitored the neurochemical profile and a robust physiological index of brain activity. Apart from studies quantifying epileptiform activity in patients (Remahl *et al*., 2008) and one Transcranial Magnetic Stimulation study (Cantello *et al*., 2007), we know of no previous experiment that tested the effects of ketogenic diets or ketone supplementation on physiological EEG activity. Using other methodologies (mainly, behavioural testing), ketogenic diets have been repeatedly associated with improved cognitive performance (Chinna-Meyyappan *et al*., 2023). For example, two acute single-dose supplementation studies reported beneficial effects of elevating βHB levels on cognitive performance in a subgroup of Alzheimer’s disease patients (Reger *et al*., 2004) and on functional network stability in healthy individuals, interpreted as a functional correlate of cognitive acuity {Mujica-Parodi, 2020 #8}. Moreover, repeated ketone supplementation was associated with improved cognitive symptoms in Mild Cognitive Impairment (Fortier *et al*., 2019).

A separate set of studies measured the effects of βHB supplementation or ketogenic diets on the concentration of GABA and glutamate (assessed invasively and/or noninvasively with MRS); however, many focused on pathological conditions: from different forms of epilepsy to neurodevelopmental disorders, or brain tumors (Machowiec *et al*., 2022). The reported outcomes are very heterogeneous; for example, elevated βHB levels have been associated with GABA increases (Dahlin *et al*., 2005; Yum *et al*., 2015; Calderon *et al*., 2017), decreases (Ciusani *et al*., 2021) and lack of change (Wang *et al*., 2003; Gzielo *et al*., 2020). Hone-Blanchett et al. (Hone-Blanchet *et al*., 2023) were the first to study the effects of a single dose βHB supplementation in healthy human participants. Using ^1^H-MRS 7T, they found a decrease of both GABA and glutamate levels in the cingulate cortices. Here we used a similar βHB supplementation approach combined with MEGA-PRESS MRS at 3T and found a significant increase of glutamate and no change of GABA+ concentrations. Despite the seeming inconsistency, we note that Hone-Blanchett et al.’s (Hone-Blanchet *et al*., 2023) effect was primarily seen in older participants, whereas the youngest individuals in their sample showed no change or even a small increment of glutamate concentration (Figure 3B of their paper), as we did. This suggests that the effects of βHB supplementation is critically dependent on age (Hone-Blanchet *et al*., 2023), and might even take opposite directions in young vs. older individuals. We speculate that an age-dependent change of glucose metabolism might be responsible for this reversal. *In-vitro* work shows that the effect of delivering βHB to cultured glutamatergic neurons is radically different depending on the co-administration of glucose. When βHB and glucose are delivered at equimolar concentrations, glutamate increases (through a shift in the equilibrium of the aspartate–glutamate aminotransferase reaction towards glutamate); however, when βHB is delivered alone, implying lower glucose availability, the effect is reversed (Lund *et al*., 2009; Lund *et al*., 2011). This might be related to glutamate being exploited to produce energy when glucose levels are critically low (Yudkoff *et al*., 2008). Based on the established knowledge that physiological aging is accompanied by glucose hypometabolism (Mosconi, 2013), we suggest that age differences in glucose metabolism may account for the opposite effects of βHB supplementation in older individuals (included in the sample of Hone-Blanchett et al.’s study) versus younger individuals (selectively tested here).

Estimates of glutamate concentration obtained with MRS pool across many compartments and functions (Rae, 2014): glutamate as neurotransmitter and glutamate used to produce energy; cytoplasmatic glutamate and glutamate stored in vesicles and glutamate released in the synaptic cleft; etc. This clearly means that MRS estimates cannot be taken as a direct index of glutamatergic signalling. However, there is evidence that glutamate concentrations estimated with MRS are systematically and positively correlated with cortical excitability, both in humans where it was indexed by TMS (Stagg *et al*., 2011), and in animal models where it was monitored by calcium imaging (Takado *et al*., 2022). In line with this, our results show a positive relationship between glutamate levels and an electrophysiological index of cortical responsivity.

One final biochemical consideration concerns the link between glutamate and GABA, the former being a precursor of the latter (Yudkoff *et al*., 2008). Increasing glutamate levels, e.g. by shifting the equilibrium of the aspartate–glutamate aminotransferase reaction, implies that more glutamate becomes accessible to the glutamate decarboxylase reaction to yield GABA. Thus, an acute increase of glutamate concentration (as we observe) could potentially lead to a secondary increase of GABA concentration over a longer time-frame (Yudkoff *et al*., 2008). Moreover, a persistent increase of βHB levels (e.g., achieved through repeated supplementation or dieting) enhances the expression of monocarboxylic acid transporters (Jensen *et al*., 2020), responsible for the passage of βHB across the blood-brain-barrier. This could enhance the βHB uptake and metabolism in neurons, and potentially promote further changes in the neurotransmitter profile. These considerations could help reconcile our observations of increased glutamate concentrations and enhanced cortical excitability following acute βHB supplementation, with the long-term beneficial effects of ketogenic diets in some forms of epilepsy, which are interpreted as resulting from a decreased cortical excitability. However, the current results offer no direct evidence on the long-term effects of βHB supplementation.

Our acute protocol produced a change of plasma βHB levels and of glucose and insulin that are compatible with previous studies (Clarke *et al*., 2012; Mikkelsen *et al*., 2015; Stubbs *et al*., 2017), including a decrement in plasma glucose. The latter is not a likely explanation for the observed changes in glutamate concentrations for three main reasons: 1) our glucose levels decreased very mildly and never reached hypoglycemia; 2) reduced glucose availability predicts a decrease of glutamate levels (Yudkoff *et al*., 2008), opposite of what we found; 3) Hone-Blanchett et al. showed that a very marked increase of glucose levels, achieved by supplementation, does not systematically alter glutamate or GABA levels as assessed with ^1^H-MRS. We did not directly measure βHB levels in the brain; however, there is clear evidence that acute βHB supplementation increases βHB levels in the brain since about 30 minutes following the oral administration (Mujica-Parodi *et al*., 2020), closely following plasma βHB concentrations.

In our result, plasma βHB levels varied little across participants, and the small differences are likely due to normal variations in gastrointestinal function, which may affect βHB hydrolysis and absorption (Clarke *et al*., 2012). As expected (Clarke *et al*., 2012), plasma βHB decayed slowly and showed only a minor change between 2 and 3h after supplementation. This rate of plasma βHB clearance was correlated with the glutamate levels measured in the visual cortex, *i.e.,* individuals with slower clearance showed higher glutamate concentrations following βHB supplementation. This correlation strengthens the link between plasma βHB concentration and glutamate concentration in the brain cortex, which in turn correlates with VEP amplitudes, closing the loop between our metabolic intervention and the observed changes in brain metabolism and function.

In conclusion, our combined EEG and MEGA-PRESS MRS results show that an acutely administered single dose of βHB supplement modulates both visual cortical responses and the neurochemical profile of the occipital cortex, consistent with increased cortical excitability. This is strong evidence for a functional shift of cortical circuits, which could help interpret the therapeutical potential of βHB supplementation.

## Limitations of the study

The first limitation our study is the small sample of participants that we had the opportunity to test. Although the a priori power analysis indicated that our sample size was sufficient to achieve the required statistical power, and although the observed effects are large (Cohen’s d ≥ 0.8), the small sample prevented us from investigating fundamental dimensions of inter-individual variability, including differences related to sex and gender.

Second, we elected to study the effects of an acute intervention. This prevents us from predicting the consequences of repeated βHB supplementation and of long-term dietary interventions, previously adopted with therapeutical aims, e.g. in epilepsy. However, we submit that understanding the transient effects of a single dose is a necessary first step towards unravelling the complex consequences that these metabolic interventions could have in pathological and physiological conditions.

## Acknowledgments

This research was funded by: the European Research Council (ERC) under the European Union’s Horizon 2020 research and innovation program grant n. 801715 (PUPILTRAITS) and n. 832813 (GenPercept); the European Union - Next Generation EU, in the context of The National Recovery and Resilience Plan, Investment 1.5 Ecosystems of Innovation, Project Tuscany Health Ecosystem (THE, CUP I53C22000780001), and of the grant PRIN 2022 (Project ‘RIGHTSTRESS—Tuning arousal for optimal perception’, Grant no. 2022CCPJ3J, CUP I53D23003960006); and by the Italian Ministry of University and Research under the program FARE-2 (grant SMILY). DM was partially supported by the Italian Ministry of Health grant RC-Linea 4-IRCCS Fondazione Stella Maris.

## Data and code availability

The current datasets will be uploaded on Zenodo before the publication of the manuscript.

## References

Achanta, L.B. & Rae, C.D. (2017) beta-Hydroxybutyrate in the Brain: One Molecule, Multiple Mechanisms. Neurochem Res, 42, 35–49.

Atallah, B.V., Bruns, W., Carandini, M. & Scanziani, M. (2012) Parvalbumin-expressing interneurons linearly transform cortical responses to visual stimuli. Neuron, 73, 159–170.

Bale, A.S., Adams, T.L., Bushnell, P.J., Shafer, T.J. & Boyes, W.K. (2005) Role of NMDA, nicotinic, and GABA receptors in the steady-state visual-evoked potential in rats. Pharmacol Biochem Behav, 82, 635–645.

Bartel, P., Blom, M., van der Meyden, C. & de Klerk, S. (1988) Effects of single doses of diazepam, chlorpromazine, imipramine and trihexyphenidyl on visual-evoked potentials. Neuropsychobiology, 20, 212–217.

Boker, T. & Heinze, H.J. (1984) Influence of diazepam on visual pattern-evoked potentials with due regard to nonstationary effects. Methodological problems. Neuropsychobiology, 11, 207–212.

Calderon, N., Betancourt, L., Hernandez, L. & Rada, P. (2017) A ketogenic diet modifies glutamate, gamma-aminobutyric acid and agmatine levels in the hippocampus of rats: A microdialysis study. Neurosci Lett, 642, 158–162.

Cantello, R., Varrasi, C., Tarletti, R., Cecchin, M., D’Andrea, F., Veggiotti, P., Bellomo, G. & Monaco, F. (2007) Ketogenic diet: electrophysiological effects on the normal human cortex. Epilepsia, 48, 1756–1763.

Carandini, M. & Heeger, D.J. (2011) Normalization as a canonical neural computation. Nat Rev Neurosci, 13, 51–62.

Chinna-Meyyappan, A., Gomes, F.A., Koning, E., Fabe, J., Breda, V. & Brietzke, E. (2023) Effects of the ketogenic diet on cognition: a systematic review. Nutr Neurosci, 26, 1258–1278.

Ciusani, E., Vasco, C., Rizzo, A., Girgenti, V., Padelli, F., Pellegatta, S., Fariselli, L., Bruzzone, M.G. & Salmaggi, A. (2021) MR-Spectroscopy and Survival in Mice with High Grade Glioma Undergoing Unrestricted Ketogenic Diet. Nutr Cancer, 73, 2315–2322.

Clarke, K., Tchabanenko, K., Pawlosky, R., Carter, E., Todd King, M., Musa-Veloso, K., Ho, M., Roberts, A., Robertson, J., Vanitallie, T.B. & Veech, R.L. (2012) Kinetics, safety and tolerability of (R)-3-hydroxybutyl (R)-3-hydroxybutyrate in healthy adult subjects. Regul Toxicol Pharmacol, 63, 401–408.

D’Andrea Meira, I., Romao, T.T., Pires do Prado, H.J., Kruger, L.T., Pires, M.E.P. & da Conceicao, P.O. (2019) Ketogenic Diet and Epilepsy: What We Know So Far. Front Neurosci, 13, 5.

Dahlin, M., Elfving, A., Ungerstedt, U. & Amark, P. (2005) The ketogenic diet influences the levels of excitatory and inhibitory amino acids in the CSF in children with refractory epilepsy. Epilepsy Res, 64, 115–125.

Erecinska, M., Nelson, D., Daikhin, Y. & Yudkoff, M. (1996) Regulation of GABA level in rat brain synaptosomes: fluxes through enzymes of the GABA shunt and effects of glutamate, calcium, and ketone bodies. J Neurochem, 67, 2325–2334.

Fischl, B., Salat, D.H., Busa, E., Albert, M., Dieterich, M., Haselgrove, C., van der Kouwe, A., Killiany, R., Kennedy, D., Klaveness, S., Montillo, A., Makris, N., Rosen, B. & Dale, A.M. (2002) Whole brain segmentation: automated labeling of neuroanatomical structures in the human brain. Neuron, 33, 341–355.

Fortier, M., Castellano, C.A., Croteau, E., Langlois, F., Bocti, C., St-Pierre, V., Vandenberghe, C., Bernier, M., Roy, M., Descoteaux, M., Whittingstall, K., Lepage, M., Turcotte, E.E., Fulop, T. & Cunnane, S.C. (2019) A ketogenic drink improves brain energy and some measures of cognition in mild cognitive impairment. Alzheimers Dement, 15, 625–634.

Garcia-Rodriguez, D. & Gimenez-Cassina, A. (2021) Ketone Bodies in the Brain Beyond Fuel Metabolism: From Excitability to Gene Expression and Cell Signaling. Front Mol Neurosci, 14, 732120.

Geller, A.M., Hudnell, H.K., Vaughn, B.V., Messenheimer, J.A. & Boyes, W.K. (2005) Epilepsy and medication effects on the pattern visual evoked potential. Doc Ophthalmol, 110, 121–131.

Gzielo, K., Janeczko, K., Weglarz, W., Jasinski, K., Klodowski, K. & Setkowicz, Z. (2020) MRI spectroscopic and tractography studies indicate consequences of long-term ketogenic diet. Brain Struct Funct, 225, 2077–2089.

Harris, A.D., Puts, N.A. & Edden, R.A. (2015) Tissue correction for GABA-edited MRS: Considerations of voxel composition, tissue segmentation, and tissue relaxations. J Magn Reson Imaging, 42, 1431–1440.

Helmholz, H.F. & Keith, H.M. (1933) TEN YEARS’ EXPERIENCE IN THE TREATMENT OF EPILEPSY WITH KETOGENIC DIET. Archives of Neurology & Psychiatry, 29, 808–812.

Hone-Blanchet, A., Antal, B., McMahon, L., Lithen, A., Smith, N.A., Stufflebeam, S., Yen, Y.F., Lin, A., Jenkins, B.G., Mujica-Parodi, L.R. & Ratai, E.M. (2023) Acute administration of ketone beta-hydroxybutyrate downregulates 7T proton magnetic resonance spectroscopy-derived levels of anterior and posterior cingulate GABA and glutamate in healthy adults. Neuropsychopharmacology, 48, 797–805.

Hudnell, H.K. & Boyes, W.K. (1991) The comparability of rat and human visual-evoked potentials. Neurosci Biobehav Rev, 15, 159–164.

Jensen, N.J., Wodschow, H.Z., Nilsson, M. & Rungby, J. (2020) Effects of Ketone Bodies on Brain Metabolism and Function in Neurodegenerative Diseases. Int J Mol Sci, 21.

Jia, K., Frangou, P., Karlaftis, V.M., Ziminski, J.J., Giorgio, J., Rideaux, R., Zamboni, E., Hodgson, V., Emir, U. & Kourtzi, Z. (2022) Neurochemical and functional interactions for improved perceptual decisions through training. J Neurophysiol, 127, 900–912.

Juge, N., Gray, J.A., Omote, H., Miyaji, T., Inoue, T., Hara, C., Uneyama, H., Edwards, R.H., Nicoll, R.A. & Moriyama, Y. (2010) Metabolic control of vesicular glutamate transport and release. Neuron, 68, 99–112.

Lin, A., Andronesi, O., Bogner, W., Choi, I.Y., Coello, E., Cudalbu, C., Juchem, C., Kemp, G.J., Kreis, R., Krssak, M., Lee, P., Maudsley, A.A., Meyerspeer, M., Mlynarik, V., Near, J., Oz, G., Peek, A.L., Puts, N.A., Ratai, E.M., Tkac, I., Mullins, P.G. & Experts’ Working Group on Reporting Standards for, M.R.S. (2021) Minimum Reporting Standards for in vivo Magnetic Resonance Spectroscopy (MRSinMRS): Experts’ consensus recommendations. NMR Biomed, 34, e4484.

Lozano-Soldevilla, D. (2018) On the Physiological Modulation and Potential Mechanisms Underlying Parieto-Occipital Alpha Oscillations. Front Comput Neurosci, 12, 23.

Lund, T.M., Obel, L.F., Risa, O. & Sonnewald, U. (2011) beta-Hydroxybutyrate is the preferred substrate for GABA and glutamate synthesis while glucose is indispensable during depolarization in cultured GABAergic neurons. Neurochem Int, 59, 309–318.

Lund, T.M., Risa, O., Sonnewald, U., Schousboe, A. & Waagepetersen, H.S. (2009) Availability of neurotransmitter glutamate is diminished when beta-hydroxybutyrate replaces glucose in cultured neurons. J Neurochem, 110, 80–91.

Machowiec, P.A., Maksymowicz, M. & Piecewicz-Szczesna, H. (2022) Magnetic resonance spectroscopy as a promising modality for assessing ketogenic diet impact on the level of cerebral metabolites in the treatment of certain neurological disorders. Ann Agric Environ Med, 29, 201–206.

Maddock, R.J. (2023) Comment on report of a large reduction in cortical GABA following ketone ingestion. Neuropsychopharmacology, 48, 847.

Mescher, M., Merkle, H., Kirsch, J., Garwood, M. & Gruetter, R. (1998) Simultaneous in vivo spectral editing and water suppression. NMR Biomed, 11, 266–272.

Mikkelsen, K.H., Seifert, T., Secher, N.H., Grondal, T. & van Hall, G. (2015) Systemic, cerebral and skeletal muscle ketone body and energy metabolism during acute hyper-D-beta-hydroxybutyratemia in post-absorptive healthy males. J Clin Endocrinol Metab, 100, 636–643.

Morrone, M.C. & Burr, D.C. (1986) Evidence for the existence and development of visual inhibition in humans. Nature, 321, 235–237.

Morrone, M.C., Burr, D.C. & Maffei, L. (1982) Functional implications of cross-orientation inhibition of cortical visual cells. I. Neurophysiological evidence. Proc R Soc Lond B Biol Sci, 216, 335–354.

Morrone, M.C., Burr, D.C. & Speed, H.D. (1987) Cross-orientation inhibition in cat is GABA mediated. Exp Brain Res, 67, 635–644.

Mosconi, L. (2013) Glucose metabolism in normal aging and Alzheimer’s disease: Methodological and physiological considerations for PET studies. Clin Transl Imaging, 1.

Mujica-Parodi, L.R., Amgalan, A., Sultan, S.F., Antal, B., Sun, X., Skiena, S., Lithen, A., Adra, N., Ratai, E.M., Weistuch, C., Govindarajan, S.T., Strey, H.H., Dill, K.A., Stufflebeam, S.M., Veech, R.L. & Clarke, K. (2020) Diet modulates brain network stability, a biomarker for brain aging, in young adults. Proc Natl Acad Sci U S A, 117, 6170–6177.

Mullins, P.G., McGonigle, D.J., O’Gorman, R.L., Puts, N.A., Vidyasagar, R., Evans, C.J., Cardiff Symposium on, M.R.S.o.G. & Edden, R.A. (2014) Current practice in the use of MEGA-PRESS spectroscopy for the detection of GABA. Neuroimage, 86, 43-52.

Newman, J.C. & Verdin, E. (2017) beta-Hydroxybutyrate: A Signaling Metabolite. Annu Rev Nutr, 37, 51–76.

Owen, O.E., Morgan, A.P., Kemp, H.G., Sullivan, J.M., Herrera, M.G. & Cahill, G.F., Jr. (1967) Brain metabolism during fasting. J Clin Invest, 46, 1589–1595.

Pan, J.W., de Graaf, R.A., Petersen, K.F., Shulman, G.I., Hetherington, H.P. & Rothman, D.L. (2002) [2,4-13 C2]-beta-Hydroxybutyrate metabolism in human brain. J Cereb Blood Flow Metab, 22, 890–898.

Poff, A.M., Rho, J.M. & D’Agostino, D.P. (2019) Ketone Administration for Seizure Disorders: History and Rationale for Ketone Esters and Metabolic Alternatives. Front Neurosci, 13, 1041.

Provencher, S.W. (2001) Automatic quantitation of localized in vivo 1H spectra with LCModel. NMR Biomed, 14, 260–264.

Puchalska, P. & Crawford, P.A. (2017) Multi-dimensional Roles of Ketone Bodies in Fuel Metabolism, Signaling, and Therapeutics. Cell Metab, 25, 262–284.

Rae, C.D. (2014) A guide to the metabolic pathways and function of metabolites observed in human brain 1H magnetic resonance spectra. Neurochem Res, 39, 1–36.

Reger, M.A., Henderson, S.T., Hale, C., Cholerton, B., Baker, L.D., Watson, G.S., Hyde, K., Chapman, D. & Craft, S. (2004) Effects of beta-hydroxybutyrate on cognition in memory-impaired adults. Neurobiol Aging, 25, 311–314.

Remahl, S., Dahlin, M.G. & Amark, P.E. (2008) Influence of the ketogenic diet on 24-hour electroencephalogram in children with epilepsy. Pediatr Neurol, 38, 38–43.

Rideaux, R. (2021) No balance between glutamate+glutamine and GABA+ in visual or motor cortices of the human brain: A magnetic resonance spectroscopy study. Neuroimage, 237, 118191.

Rideaux, R., Ehrhardt, S.E., Wards, Y., Filmer, H.L., Jin, J., Deelchand, D.K., Marjanska, M., Mattingley, J.B. & Dux, P.E. (2022) On the relationship between GABA+ and glutamate across the brain. Neuroimage, 257, 119273.

Sanaei Nezhad, F., Anton, A., Michou, E., Jung, J., Parkes, L.M. & Williams, S.R. (2018) Quantification of GABA, glutamate and glutamine in a single measurement at 3 T using GABA-edited MEGA-PRESS. NMR Biomed, 31.

Smith, M.A., Bair, W. & Movshon, J.A. (2006) Dynamics of suppression in macaque primary visual cortex. J Neurosci, 26, 4826–4834.

Soto-Mota, A., Norwitz, N.G. & Clarke, K. (2020) Why a d-beta-hydroxybutyrate monoester? Biochem Soc Trans, 48, 51–59.

Stagg, C.J., Bestmann, S., Constantinescu, A.O., Moreno, L.M., Allman, C., Mekle, R., Woolrich, M., Near, J., Johansen-Berg, H. & Rothwell, J.C. (2011) Relationship between physiological measures of excitability and levels of glutamate and GABA in the human motor cortex. J Physiol, 589, 5845–5855.

Steel, A., Mikkelsen, M., Edden, R.A.E. & Robertson, C.E. (2020) Regional balance between glutamate+glutamine and GABA+ in the resting human brain. Neuroimage, 220, 117112.

Stubbs, B.J., Cox, P.J., Evans, R.D., Santer, P., Miller, J.J., Faull, O.K., Magor-Elliott, S., Hiyama, S., Stirling, M. & Clarke, K. (2017) On the Metabolism of Exogenous Ketones in Humans. Front Physiol, 8, 848.

Takado, Y., Takuwa, H., Sampei, K., Urushihata, T., Takahashi, M., Shimojo, M., Uchida, S., Nitta, N., Shibata, S., Nagashima, K., Ochi, Y., Ono, M., Maeda, J., Tomita, Y., Sahara, N., Near, J., Aoki, I., Shibata, K. & Higuchi, M. (2022) MRS-measured glutamate versus GABA reflects excitatory versus inhibitory neural activities in awake mice. J Cereb Blood Flow Metab, 42, 197–212.

Thiele, E.A. (2003) Assessing the efficacy of antiepileptic treatments: the ketogenic diet. Epilepsia, 44 **Suppl 7**, 26–29.

van Kerkoerle, T., Self, M.W., Dagnino, B., Gariel-Mathis, M.A., Poort, J., van der Togt, C. & Roelfsema, P.R. (2014) Alpha and gamma oscillations characterize feedback and feedforward processing in monkey visual cortex. Proc Natl Acad Sci U S A, 111, 14332–14341.

Wang, Z.J., Bergqvist, C., Hunter, J.V., Jin, D., Wang, D.J., Wehrli, S. & Zimmerman, R.A. (2003) In vivo measurement of brain metabolites using two-dimensional double-quantum MR spectroscopy--exploration of GABA levels in a ketogenic diet. Magn Reson Med, 49, 615–619.

Yudkoff, M., Daikhin, Y., Horyn, O., Nissim, I. & Nissim, I. (2008) Ketosis and brain handling of glutamate, glutamine, and GABA. Epilepsia, 49 **Suppl 8**, 73–75.

Yudkoff, M., Daikhin, Y., Melo, T.M., Nissim, I., Sonnewald, U. & Nissim, I. (2007) The ketogenic diet and brain metabolism of amino acids: relationship to the anticonvulsant effect. Annu Rev Nutr, 27, 415–430.

Yum, M.S., Lee, M., Woo, D.C., Kim, D.W., Ko, T.S. & Velisek, L. (2015) beta-Hydroxybutyrate attenuates NMDA-induced spasms in rats with evidence of neuronal stabilization on MR spectroscopy. Epilepsy Res, 117, 125–132.

