## Supplementary material for "Acute supplementation of beta-hydroxybutyrate increases visual cortical excitability in humans: a combined EEG and MRS study"

### Supplementary Tables

**Table S1:** **Minimum Reporting Standards in MRS checklist**

| Site (Name or Number) | Foundation CNR-Regione Toscana G.Monasterio |
| --- | --- |
| 1. Hardware |  |
| a. Field strength [T] | 3 |
| b. Manufacturer | GE – General Electric |
| c. Model | GE HDx TWINSPEE |
| d. RF coils | 8-channel receive head coil |
| e. Additional hardware | N/A |
| 2. Acquisition |  |
| a. Pulse sequence | MEGA-PRESS |
| b. Volume of Interest (VOI) locations | Visual cortex (centered on right calcarine sulcus) |
| c. Nominal VOI size [cm^3^, mm^3^] | 25x25x25 mm |
| d. Repetition Time (TR), Echo Time (TE) [ms, s] | TR=1500, TE=68ms |
| e. Total number of Excitations or acquisitions per spectrum | 320 |
| f. Additional sequence parameters: | Spectral bandwidth: 5000 Hz  Spectral points: 4096 |
| g. Water Suppression Method | Standard |
| h. Shimming Method, reference peak, and thresholds for “acceptance of shim” chosen | Automatic 3D head shim to achieve water peak linewidth below 15 Hz. |
| i. Triggering or motion correction method | N/A |
| 3. Data analysis methods and outputs |  |
| a. Analysis software | MRspa (preprocessing, version v1.5c), LCModel (fitting and quantification) |
| b. Processing steps deviating from quoted reference or product | MRspa pre-processing options selected:  - eddy current corr.: ECC2 + zero phase  - frequency corr.: absolute (3.01)  - phase corr.: least square |
| c. Output measure | Tissue-corrected concentrations relative to water or NAA |
| d. Quantification references and assumptions, fitting model assumptions | We fitted model spectra of glutamate (Glu), glutamine (Gln), γ-amino-butyric acid (GABA), glutathione (GSH) and N acetylaspartate (NAA) to the edited spectra.  We used the dedicated MEGA-PRESS basis set (‘mega-press 3’) on LCModel. |
| 4. Data Quality |  |
| a. Reported variables | See Table S2 |
| b. Data exclusion criteria | Water peak linewidth > 15 Hz  CRLB > 10%  Lipid contamination by visual inspection |
| c. Quality measures of postprocessing Model fitting | See Table S2 |
| d. Sample Spectrum | See Figure 3B for average across- participants |

**Table S2. MRS metabolite concentrations and quality measures:** Glutamate and GABA+ values before alpha-correction and referenced on water, Cramer-Rao lower bound (CRLB), water peak linewidth and signal-to-noise ratio (SNR) are shown for the two MRS voxels (pre and post βHB supplementation).

| MRS voxel | MRS quality measure | Mean | Std |
| --- | --- | --- | --- |
| Early Visual  Pre βHB | Glu | 0.82 | 0.20 |
|  | GABA+ | 0.78 | 0.06 |
|  | Glu CRLB | 7.50 | 1.64 |
|  | GABA+ CRLB | 3.30 | 0.48 |
|  | Linewidth | 11.03 | 1.55 |
|  | SNR | 23.60 | 3.09 |
| Early Visual  Post βHB | Glu | 0.92 | 0.18 |
|  | GABA+ | 0.78 | 0.06 |
|  | Glu CRLB | 5.50 | 0.52 |
|  | GABA+ CRLB | 3.00 | 0.00 |
|  | Linewidth | 10.37 | 0.75 |
|  | SNR | 28.50 | 3.72 |
